# Investigating the role of sensorimotor spatial dependencies in shaping conscious access to virtual 3D objects

**DOI:** 10.1101/2024.12.30.630721

**Authors:** Paweł Motyka, David J. Schwartzman, Anil K. Seth, Keisuke Suzuki

## Abstract

According to sensorimotor accounts of perceptual experience, the subjective veridicality of an object (the sense of its ‘presence’) builds up gradually as one learns how changes in sensory inputs depend on bodily movements. To investigate how sensorimotor interactions shape visual experience, we designed a virtual-reality-based study that allowed us to manipulate the complexity of spatial dependencies governing interactions with unfamiliar 3D objects. Participants had to learn to manually control fully visible objects that could move in congruent, opposite, novel (orthogonal), or random directions in response to their movements. The sensorimotor control tasks occurred alternately with a continuous flash suppression (CFS) task evaluating the access of stationary objects to visual awareness, operationalised as the time taken for a 3D object to break the interocular suppression. We hypothesised that objects whose motion was experienced as depending on actions in a lawful, and thus encodable, manner (i.e., according to a congruent, opposite, or novel – but not random – dependency) would overcome suppression faster than objects moving randomly in response to actions (for which there is no world-related statistical structure to learn). However, while performance in the sensorimotor tasks consistently decreased along with the difficulty of the conditions (i.e., congruent > opposite > novel > random), the pre-registered analysis yielded no significant differences in breakthrough times of objects manipulated under different coupling rules. An exploratory analysis assessing whether the acquisition of ‘sensorimotor mastery’ was associated with reduced breakthrough times also revealed no significant effects. Thus, our results suggest that one’s knowledge of how an object responds to action does not play a salient role in determining conscious access to visual stimuli. This extends previous evidence for a general ineffectiveness of sensorimotor spatial manipulations in interocular suppression paradigms. Notably, in all conditions, object movement remained tightly coupled (i.e., contingent) to the participant’s actions – and given such stimuli have already been shown to break suppression faster than uncoupled/pre-recorded visual inputs – it is possible that sensorimotor contingency was a sufficiently salient factor to override any influences related to how identifiable specific spatial dependencies were.

**Highlights:** Learning sensorimotor control over objects was examined in novel VR tasks.

We varied the complexity of spatial dependencies between manual and visual rotations.

Controllable objects were expected to enter visual awareness faster in b-CFS.

No differences in breakthrough times for differently complex spatial coupling rules.

Sensorimotor contingency outweighs spatial congruence in affecting visual awareness.

## 1. Introduction

Closed loops between actions and sensory feedback provide information about the structure of the environment and the scope of possible interactions with surrounding objects. The identification of sensorimotor dependencies has been proposed to play a key role in shaping the characteristics of perceptual experience. According to sensorimotor accounts of perceptual experience (Noë, 2004; O’Regan & Noë, 2001; Seth, 2014a), subjective veridicality/presence of objects builds up as one learns how changes in incoming sensory inputs depend on bodily movements, such as of the eyes, head, or hands. Building upon the predictive processing framework (Clark, 2013; Hohwy, 2013; Seth, 2014b), it has been postulated that the degree of perceptual presence (quality of “objecthood”) corresponds to the counterfactual richness of the neural model encoding expected changes in sensory signals under a broad repertoire of possible – not necessarily executed – actions (Seth, 2014a). In a nutshell, whereas counterfactually-rich models are thought to underlie the perception of natural objects – for which there is a complex world-related statistical structure to learn (and thus a variety of sensorimotor dependencies can be inferred) – counterfactually-poor models may underlie subjectively non-veridical perceptual phenomena, such as synaesthetic concurrents or hallucinations, that do not correspond to encodable external structures. While different theoretical accounts diverge on whether the process of acquiring so-called sensorimotor mastery can be operationalised in representational terms (O’Regan & Noë, 2001; Seth, 2014a), they agree on the principle that visual experience is actively shaped by knowledge of how consistently one’s movements translate into changes in visual input.

Given that the perception of the external world typically aligns with the lifelong-learned sensorimotor dependencies, their disruption within an experimental context provides a means of uncovering their role in shaping perceptual experience. For example, while wearing visual field inverting glasses, normal visual experience has been shown to be temporarily distorted until re-adaptation to the unusual relationships between head movements and accompanying changes in visual perspective occurs (Degenaar, 2014; Kohler, 1964; Sachse et al., 2017). The factors that lead to perceptual presence, including sensorimotor dependencies, have also been extensively explored in research on sensory substitution (e.g. Auvray et al., 2007; Bermejo et al., 2015) and sensory augmentation (Kaspar et al., 2014; Schumann & O’Regan, 2017), in which newly experienced patterns of sensory stimulation have been shown to gain perceptual presence following action-based training. While research on sensorimotor dependencies has focused predominantly on the perception of suprathreshold (i.e. already accessible) stimuli, interocular suppression paradigms – such as binocular rivalry (Alais & Blake, 2005) and continuous flash suppression (CFS; Stein, 2019; Tsuchiya & Koch, 2004) – allow the examination of whether visual inputs that systematically depend on one’s actions gain preferential access to perceptual awareness. According to the predictive processing-based account of interocular rivalry (Hohwy et al., 2008; Jack & Hacker, 2014; Parr et al., 2019), conscious access to a given stimulus reflects the brain’s current best guess about the environmental causes of the incoming sensory signals. Simply put, if competing images are balanced with respect to their low-level features, the perceptual prioritisation of one stimulus over another could be interpreted as due to it being inferred as a veridically present part of the environment. While such prioritisation can be driven by consistency with signals from the other senses (e.g., Hense et al., 2019; Lunghi et al., 2014; Parker & Alais, 2006), the evidence for whether action-dependent visual inputs show preferential access to awareness is mixed.

Importantly, sensorimotor dependencies and manipulations thereof can take gdifferent forms. We use the term “sensorimotor dependency” as the most general term for any action-perception association; we use “congruency” to denote a particular form of spatial dependency, when sensory consequences follow actions in expected ways, and we use “contingency” to imply close temporal coupling between action and sensory consequences. Among these forms, *sensorimotor contingency* has been consistently reported to affect visual awareness (Suzuki et al., 2019). It has been shown that stimuli whose movement is coupled to the dynamics of a person’s movements show enhanced access to visual awareness as compared to uncoupled/pre-recorded stimuli – taking the form of 2D spheres presented during binocular rivalry (Maruya et al., 2007) or 3D virtual objects viewed in the CFS paradigm (Suzuki et al., 2019; see also Veto et al., 2018). In a similar vein, immersive visualisations of self-motion that were consistent with walking speed are prioritised during binocular rivalry compared to non-veridically fast or slow visualisations (Motyka et al., 2021a). Notably, the presence of sensorimotor *contingency* does not entail that action-induced sensory changes have to be *congruent* with expectations about how exactly the stimulation should change. A typically explored form of *sensorimotor congruency* is the spatial correspondence between the direction of one’s actions and the direction of visual movement (e.g., of an object or optic flow patterns). Interestingly, no evidence for perceptual prioritisation was found for stimuli that were spatially congruent with manual movements (Dogge et al., 2018; Suzuki et al., 2019) or active locomotion (Paris, 2017; Motyka et al., 2021b) as compared to stimuli that violated spatially-defined expectations. While it is possible that sensorimotor contingency may be a much more salient factor in determining perceptual access to visual inputs – as it signals the presence of some form of causal relationship – the apparent irrelevance of sensorimotor spatial dependencies in the wider theoretical context of this field remains puzzling.

The present study aims to address this issue from a new angle by assessing whether the mere identifiability of sensorimotor spatial coupling rules facilitates perceptual awareness of visual objects. Importantly, prior studies in this domain relied on dependencies that were either congruent or opposite to lifelong expectations (e.g., left manual rotations yielding leftward or rightward visual shifts, respectively; cf. Suzuki et al., 2019; Dogge et al., 2018). While such conditions allowed the investigation of effects attributable to pre-existing sensorimotor knowledge, on a more abstract level, both congruent and opposite action-perception mappings could be considered equivalent with respect to the complexity of the underlying sensorimotor rules and their repeatability over the experimental sessions. This might be important given the capacity of a cognitive system to rapidly identify and adapt to novel (e.g., context-specific) sensorimotor dependencies (Held, 1965; Wähnert & Gerhards, 2022; Yamamoto et al., 2006). For example, the perceived rotation of an ambiguous (suprathreshold) motion stimulus has been shown to be differentially affected by manual movements performed using a belt or cogwheel-based device (operating according to congruent and opposite action-effect couplings, respectively; Veto, Uhlig et al., 2018). It is possible that once the rules governing interactions with given objects are identified – and turn out to be equally complex and situationally persistent – the resultant subjective veridicality of these objects would also be comparable. To put it in theoretical terms (Seth, 2014a), the underlying neural models encoding the scenarios of possible interactions with consistently incongruent objects could end up being equally rich/informative as the models encoding consistently congruent responses to one’s actions. This might explain the ineffectiveness of sensorimotor congruency manipulations in interocular suppression paradigms (Dogge et al., 2018; Suzuki et al., 2019), which, overall, contrasts with the evidence of decreased perceptual access to uncoupled stimuli, for which – due to the lack of both temporal and spatial correspondence to motor movements (Maruya et al., 2007; Suzuki et al., 2019) – building a reliable model is rendered impossible.

To investigate the relevance of the ability to encode how visual stimulation changes with bodily movements, we developed novel *sensorimotor mastery tasks* in which interactions with unfamiliar 3D objects – observed in an immersive virtual reality – could unfold according to spatial dependencies of differing complexities. Throughout the tasks, participants had to manually control a fully visible object to reach a visually cued static rotation, or to follow as closely as possible a trajectory of continually shifting rotation. In different experimental conditions, virtual objects were observed to rotate in either congruent, opposite, novel (i.e., orthogonal) or – critically – random directions in response to manual rotations performed using a trackball device. These goal-oriented tasks were used to assess the participants’ performance and allowed for prolonged exposure to associations between distinctive visual objects and different coupling rules. Throughout the experimental session, sensorimotor mastery tasks were presented alternately with the CFS blocks, during which the previously controlled objects underwent perceptual suppression.

We hypothesised that virtual objects whose motion was experienced as depending on actions in a lawful, and thus encodable, manner (i.e., according to a congruent, opposite, or novel spatial dependency) would break through suppression faster than objects that shift randomly in response to actions (for which there is no world-related statistical structure to learn). Thus, the level of conscious access to visual inputs was expected to be mediated by the *identifiability* of the rules governing sensorimotor interactions, which is typical for real-world objects, but less evident or absent in the case of subjectively unreal percepts (e.g., hallucinations). Additionally, we aimed to explore whether perceptual performance improved with progress in the acquisition of *sensorimotor mastery*, operationalised as the ability to control the objects’ movement. This exploration pertained specifically to the opposite and novel sensorimotor conditions, where actual learning over time was expected – as opposed to the congruent and random conditions, where near-perfect and chance-level performances (respectively) were anticipated. The study design and planned analyses were pre-registered (https://osf.io/mqvj2).

## 2. Material and methods

### 2.1. Stimuli

All four visual objects presented throughout the study were created using Blender (The Blender Foundation, Amsterdam, Netherlands). They were designed to be unfamiliar but distinguishable abstract 3D structures with a roughly spherical shape, but also to display some irregularities that allow their rotation to be clearly discriminated. The objective was to create objects that were similar with respect to basic characteristics (such as size: 12 x 12 x 12 cm, spherical shape, colour, and complexity), but at the same time distinctive enough to be easily distinguishable from each other (Fig. 1a). Pilot participants were able to correctly identify the presented objects in 98.0% of trials, based on reported detection. The suppression durations for the created stimuli (when presented under b-CFS conditions) were pre-tested in a pilot study (n = 10) using a one-way repeated-measures analysis of variance (ANOVA). The post-hoc Bonferroni-corrected comparison indicated no significant differences in suppression time between objects 1, 2, and 4 (*p* values range: 0.147–0.630). However, object 3 – in its original form – was observed to be detected significantly later than the other objects (*p* values range: 0.001–0.024; *F*(3, 27) = 5.97, *p* = 0.016, ε = 0.55, η^2^_*G*_ = 0.130; GG-corrected), so it underwent slight alterations to increase its visual complexity (see Figure 1). A follow-up pilot test with three observers yielded more similar raw detection times for the modified Object 3 and other objects (with average detection times: Obj. 1 = 3.50 s, Obj. 2 = 3.26 s; Obj 3 = 3.56 s; Obj. 4 = 3.72 s). Based on these results, we decided to use this set of objects in the final study (which by design did not require equal detection times due to a fully counterbalanced conditions; see Sections 2.4 and 2.5).

**Figure 1.**
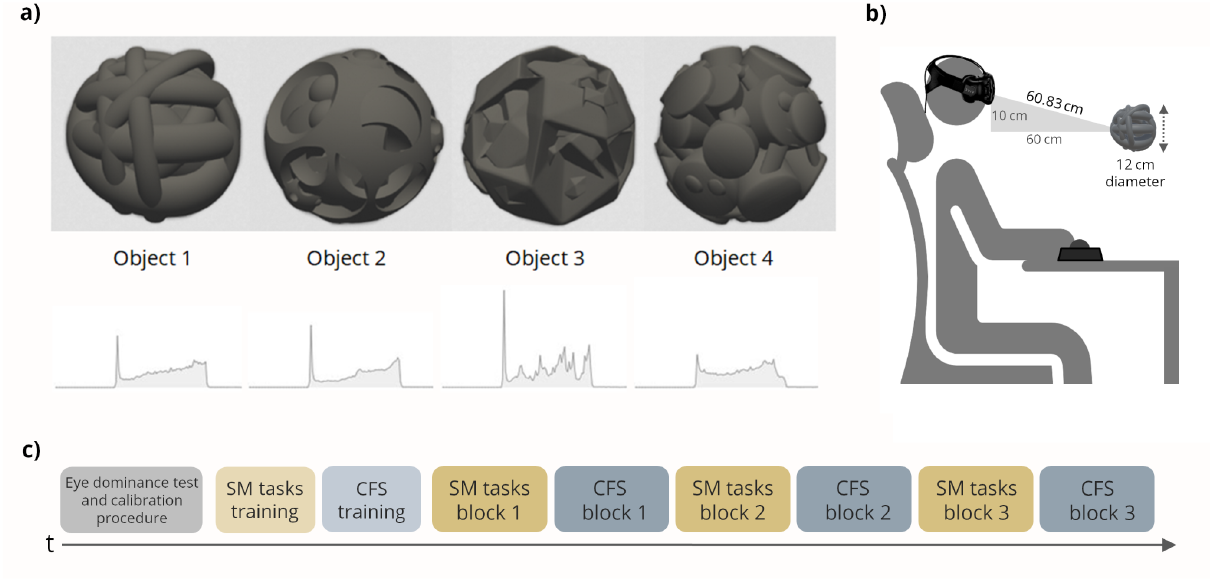
Experimental setup. **(a)** The virtual 3D objects presented during the tasks and their respective luminosity histograms. **(b)** A schematic illustration of a virtual-reality-based experimental setup. **(c)** The experiment consisted of alternating blocks of newly designed sensorimotor mastery tasks (see Figure 2) and an adapted version of the continuous flash suppression paradigm (see Figure 3).

### 2.2. Setup and tasks

During the experiment, participants were seated at a desk and wore a head-mounted virtual reality (VR) display (HTC Vive, HTC, Taiwan) while keeping their non-dominant hand on the keyboard and the dominant hand on the trackball device (Kensington Trackball Expert). The VR environment and experimental procedure were developed using Unity version: 2018.2.11f (Unity Technologies, San Francisco, CA). Before the start, participants were asked to adopt a comfortable position that they can maintain during the experiment. The position of the head served as the default camera (i.e., first-person viewpoint) position in the virtual scene. Throughout the study, participants were presented with a virtual room in which task-related objects appeared in front of them within a reachable distance, slightly below eye level (i.e., the camera is located 10 cm up from and 60.83 cm away from the centre of the objects; Fig. 1b). If the head position deviated more than 8 cm in any direction from the initial setup, a warning beep was played, and the participants were asked to return to the original head position. The fixed frame rate within Unity was 90 frames per second. In summary, the procedure consisted of interchangeable blocks of the CFS and “Sensorimotor Mastery” (SM) tasks, preceded by the eye dominance test (Miles, 1930) and the calibration procedure accounting for individual differences in the pupillary distance (Fig. 1c).

### 2.3. Sensorimotor Mastery Tasks

The exposure to different sensorimotor spatial dependencies was embedded in goal-directed tasks allowing the assessment of participants’ ability to control the movement of objects under differently complex coupling rules. Individual trials entailed the presentation of a fully visible virtual object, whose rotation was changing simultaneously with manual actions performed using a trackball device (along the *x*- and *y*-axes). Crucially, the direction of action-induced visual rotation varied depending on the sensorimotor conditions (referred to as: “congruent”, “opposite”, “novel”, and “random”; Fig. 2a). In the congruent condition, the direction of observed rotations simply matched the direction of manual movements. The opposite condition was the inverse of the congruent condition, that is leftward rotations yielded rightward visual shifts (and vice versa) and upward rotations yielded downward visual shifts (and vice versa). In the “novel” condition, visual consequences were neither congruent nor opposite to manual actions: horizontal movements evoked rotation along the vertical axis (according to left-up and right-down associations), while vertical movements induced horizontal shifts (according to up-right and down-left associations). Finally, the random condition operated in a more complex way as the coupling rule switched with each newly initiated action, that is when there was at least a 200 ms pause since the last manual movement, or when there was a substantial change in movement direction – exceeding a threshold of 60° relative to the direction of the previous action. At the moment of the switch, one of eight possible coupling rules was randomly selected – there were four associations in which vertical/horizontal actions yielded vertical/horizontal rotations and four associations in which vertical/horizontal actions produced horizontal/vertical rotations, respectively (hence, all possible combinations were utilised). The random condition was designed in order to give the feeling that visual changes, despite being *contingent* on one’s actions, were not spatially consistent or predictable in the sense that they did not play out according to any identifiable coupling rule – making it impossible to voluntarily steer an object’s movement. Each participant was exposed to all sensorimotor conditions. The assignment of particular visual objects (1, 2, 3, 4; Fig. 1) to sensorimotor conditions was constant throughout the experiment for a given participant, but it was counterbalanced between participants to satisfy all possible combinations (i.e., 24 unique object-condition pairings; n = 48, see section *Participants*).

**Figure 2.**
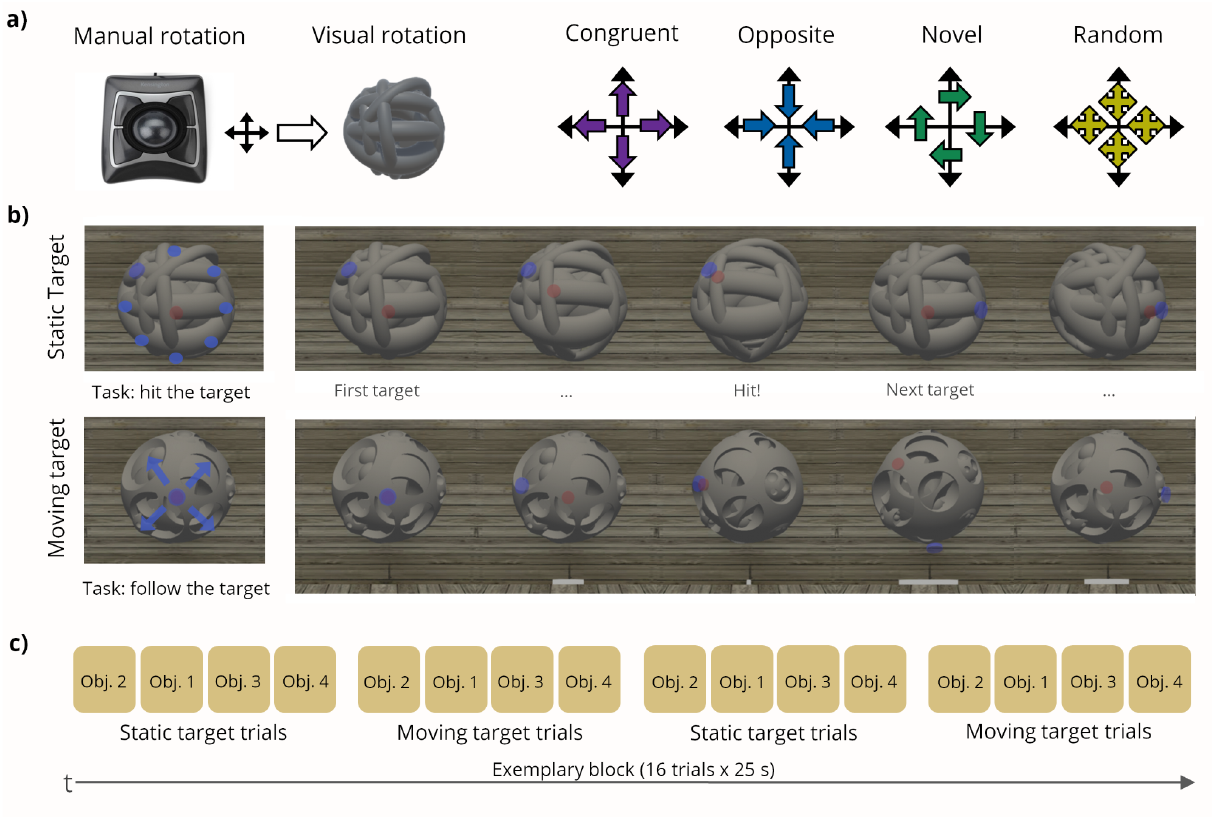
Sensorimotor mastery tasks were designed to assess participants’ ability to control the movement of virtual objects under different sensorimotor spatial dependencies. **(a)** A schematic depiction of the condition-specific mappings between manual manipulations of a trackball device and concomitant rotation of virtual objects. **(b)** In the static target task, participants had to rotate an object to reach visually cued stationary targets (blue cylinders) with a red pointer attached to the rotated object. In the moving target task, they had to rotate an object to follow as closely as possible the continuously shifting target. **(c)** During a single block, the two tasks were presented alternately. Each participant was exposed to all four sensorimotor conditions – their assignment to particular visual objects was fixed for a given participant but counterbalanced across participants.

Overall, there were two interchangeably presented versions of the task differing in what type of control over the object was required from the participants. In the “static target” task (Fig. 2b; supplementary video 1), the goal was to rotate a virtual object so as to reach a visually cued stationary target (represented by a semi-transparent blue flat cylinder) with a semi-transparent red pointer, that moved along with the rotated figure. After reaching the target, the object’s rotation returned to the default position and the new target appeared in one of eight possible positions (distributed as if around a clock dial). Subsequent static target positions were drawn randomly from an 8-item list (with no replacement), with a new list being reinstated when all items were exhausted (so to reduce the chances of presenting the same target item twice in a row). The overarching goal was to reach as many targets as possible during a single trial lasting 25 seconds. In each trial, only one virtual object (and thus one sensorimotor coupling rule) was present. Correspondingly, in the “moving target” task (Fig. 2c; supplementary video 2), the participant’s goal was to rotate a virtual object to follow a continuously shifting rotation cued by a blue target. The target moved in smooth transitions between extreme positions located on the surface of the invisible sphere (as if around a “clock dial”). After reaching a given extreme position, the target started to move in the opposite direction with some randomness (from 0 to 60°) in how much its trajectory deviated from a direct path towards the opposite position (making its movement partly but not fully predictable). The distance between the red pointer and the target was visualised in real-time in the form of a grey bar of changing length. The participant’s goal was to follow the indicated rotation as closely as possible (i.e., to keep the distance indicating bar as short as possible). As in the static target task, a single trial lasted 25 seconds and entailed the presentation of only one of the virtual objects (and thus one sensorimotor coupling rule).

In both static and moving target tasks, individual trials were organised into segments of four trials featuring four different objects. The procedure has been designed this way so as not to switch the task type every trial, but only between segments. The entire block consisted of sixteen trials (4 segments x 4 trials, Fig. 2d; amounting to 6 minutes and 40 seconds of active control of the objects per block). The practice block was respectively shorter (2 segments x 4 trials). Trials were separated by the intertrial breaks terminated upon a space bar keypress. The order of objects within each segment was fixed throughout a given block but was randomised in each new block so to prevent the same objects (and thus sensorimotor conditions) from being repeated one after another. For standardisation purposes, in both tasks, an excessively slow or fast manual rotation (i.e., below 0.25 or above 3 degrees/frame) triggered the display of respective warnings stating the need to increase or reduce the speed of action. Rotation speeds were calculated as a rolling average across a 3000-ms time window. Before the start of the practice block, participants were informed that these tasks might sometimes seem easy and sometimes hard, but that they should try to do their best regardless. After completing all blocks, they were debriefed that for one of the objects it was not possible to learn how to control it (i.e., that it behaved randomly in response actions).

### 2.4. Continuous Flash Suppression

During the CFS task, virtual objects were presented via the head-mounted display to the nondominant eye, while a dynamic Mondrian mask was shown in the dominant eye (Fig. 3; supplementary video 3). The Mondrian mask (22 x 22 cm) was horizontally and vertically centred on the virtual object, but positioned 3 cm closer to the observers relative to the centre of the object. The mask consisted of 400 rectangles of varying size, colour, and movement direction. The height and width of the rectangles were randomly selected varying from approximately 1 mm to 4 cm. Each rectangle was randomly assigned one of 10 possible colours and moved in one of 8 possible directions (along the horizontal, vertical, and diagonal axes) at a constant speed of 2 cm/s (for horizontal/vertical directions) or 6.14 cm/s (for diagonal directions). A grey cross was presented in the centre of the mask as a fixation point, to help minimise eye movements during the experiment. Participants were also asked to keep their gaze fixed on the cross in the centre of the mask and to avoid blinking during the first phase of the trial when the mask is visible. In each trial, a stationary object appeared between 300 and 600 ms after the start of the trial and gradually increased its opacity from 0% to 100% in the interval of 2000 ms. The vertical position of the object was jittered randomly in the small range of −0.5 to 0.5 cm relative to the default position (0 cm), in order to make it difficult to look out for specific features of the objects in repetitive locations. Participants were instructed to report the appearance of the object with the space bar (1st response: *detection*). This terminated the stimulus presentation and evoked the scene with all objects, from which participants had to choose the one they had seen (2nd response: *identification;* 4-AFC task). The objects were allocated randomly across the four quadrants and participants had to report the location of the correct object by pressing one of the four trackball keys (upper-right, upper-left, bottom-right, bottom-left). If there was no object detection within a predefined window of 7000 ms, the presentation ended automatically, and participants were encouraged to guess which object might have been shown. Participants were instructed to report the detection of an object as quickly as possible and later identify it focusing on accuracy rather than speed (for a similar approach see Zopf et al., 2019). The two-stage response was favoured in order to isolate the potential confounds associated with choosing between four buttons from the detection times themselves. The CFS trials were separated by the 500 ms intertrial intervals in which an empty room was presented.

**Figure 3.**
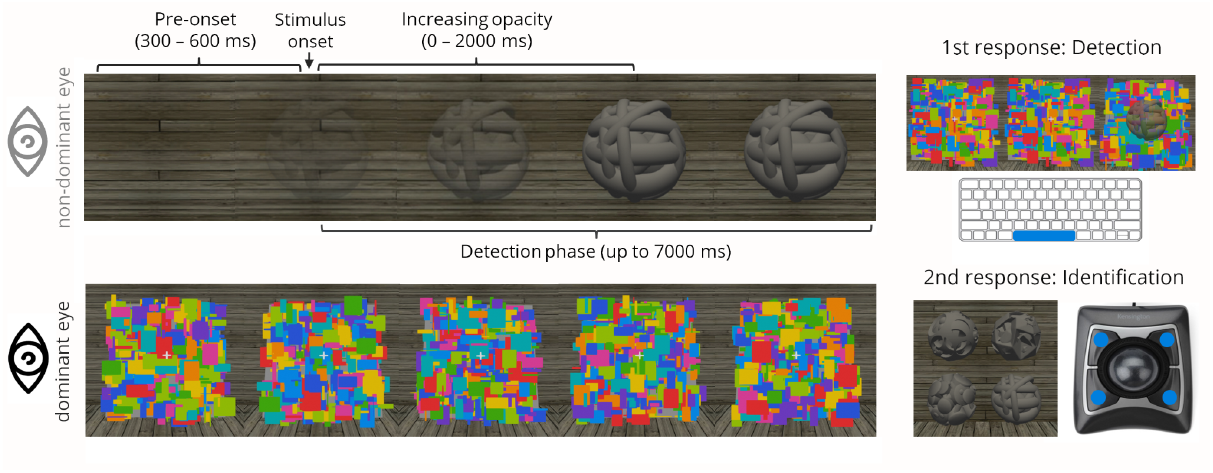
Schematic depiction of an experimental trial in the continuous flash suppression (CFS) task. The dynamic Mondrian mask was presented in the dominant eye and a virtual object in the nondominant eye via the head-mounted display. Participants had to (1) report immediately the detection of an object by pressing the spacebar and subsequently (2) choose which of four objects had been seen by pressing one of the four trackball keys.

There were three experimental blocks, each consisting of 52 trials – 13 trials per object. The trials were presented in pseudo-random order set at the start of each block. The practice block mirrored the structure of the experimental blocks but was respectively shorter (24 trials; 6 trials per object). Participants were encouraged to take a short break after completing a full CFS block. The unity project with the procedure codes can be shared upon request.

### 2.5. Participants

Participants with normal or corrected-to-normal vision and no history of amblyopia, colour-blindness, psychiatric or neurological conditions were recruited to take part in the study. As stated in the preregistration (https://osf.io/mqvj2), we aimed to acquire data from 48 participants. The choice of sample size was based both on design-specific constraints and statistical power considerations. There were 24 possible assignments of particular objects (1, 2, 3, 4) and sensorimotor conditions (congruent, opposite, novel, random), hence, to obtain a fully counterbalanced design, our choice was practically limited to multiples of 24. Given the novelty of the hypotheses tested and the lack of prior analogous studies, we set out to obtain a sample size that should allow for detecting medium size effects (i.e., Cohen’s *d* = 0.5) with a power of at least 80% in each of the three Bonferroni-corrected comparisons of interest (alpha = 0.017, two-tailed tests; see *Data analysis*). An analysis performed using G*power software indicated that a sample size of 48 would provide sufficient power (i.e., 83%) in each of these comparisons, thus this sample size was targeted. Notably, it exceeded the post-exclusions sample sizes used in the most similar of the previous CFS studies: Suzuki et al., 2019: n = 29 (exp. 1), n = 29 (exp. 2); Korisky & Mudrik, 2021: n = 17 (exp. 1), n = 15 (exp. 2), n = 33 (exp. 3). Altogether 52 participants were recruited, of whom 1 was excluded due to the excessive speed of manual movements (see *Exclusion criteria*) and 3 discontinued the session after experiencing no perceptual breakthroughs in the CFS practice block. Data from a counterbalanced sample of 48 participants (28 females, mean age = 23.1, *SD* = 3.78, range: 18–32 years) were subjected to final analysis. All participants gave written informed consent before taking part in the study and were financially compensated for their participation. The procedure was approved by the Ethics committee of the Faculty of Psychology at the University of Warsaw.

### 2.6. Data preprocessing

The acquired data were pre-processed and analysed according to the pre-registered plan (https://osf.io/mqvj2). First, all CFS trials with incorrect identification of the reportedly detected stimulus were removed from the main analysis. Then, for each participant, median detection times were derived for each of the four sensorimotor conditions. The use of medians was preferred to allow taking into account trials with no detection within the predefined 7000-ms time window (indicating the longest suppression durations). The removal of these – nevertheless informative – trials would result in including more data in the conditions that yielded faster detections, thus undermining a fair comparison between the conditions (for more on this approach, see Gayet et al., 2016; Gayet & Stein, 2017). In the sensorimotor mastery tasks, performance measures were operationalised in two ways. In the static target task, the mean number of hits during the trials (i.e., instances of reaching the cued target by the pointer) was derived separately for each participant and condition. In the moving target task, the median distance between the target and the pointer throughout the trials was derived in an analogous manner. In this case, the use of medians was favoured in order to minimise the impact of the occasional intervals with a large distance of the pointer to the target (logged on a frame-by-frame basis) on the resultant averages.

### 2.7. Exclusion criteria

Data from participants were excluded prior to the analysis if at least one of the pre-registered criteria for exclusion was met. (1) More than 45% of trials with no *detection* (i.e., no perceptual breakthrough) in any of the four conditions (i.e., particular objects). It was assumed that meeting this criterion would indicate extreme perceptual suppression impinging on the calculation of median detection times for a given participant and condition. (2) More than 15% of trials with incorrect *identification* of the reportedly detected objects. It was assumed that meeting this criterion would suggest that the participant did not pay sufficient attention during the task. (3) Median detection time lower than 500 ms (all trials considered). It was assumed that meeting this criterion would indicate unusually weak or no genuine perceptual suppression. (4) Excessively slow or fast velocity of the manual actions during the sensorimotor mastery tasks defined as the average speed outside the expected range signalled by the display of warnings (i.e., lower than 0.25 degrees/frame or higher than 3 degrees/frame). (5) No better performance in the congruent condition as compared to other conditions (taken together) during the sensorimotor mastery tasks. It was assumed that meeting these criteria would suggest that the participant did not pay sufficient attention during the tasks. Altogether, only one participant was excluded based on these criteria (criterion 4; see *Participants*).

### 2.8. Hypothesis testing

It was hypothesised that virtual objects whose motion was experienced as depending on the participant’s actions in a lawful, and thus encodable, manner (i.e., according to a congruent, opposite, or novel dependency) would be detected faster than objects moving randomly in response to actions. A one-way repeated-measures analysis of variance (ANOVA) was used to compare breakthrough times for objects manipulated under different sensorimotor conditions. In addition to a significant main effect for the condition, the following results of the three planned Bonferroni-corrected two-tailed comparisons were anticipated: in the congruent (1), opposite (2), and novel (3) conditions, breakthrough times would be faster as compared to the random condition. The Greenhouse–Geisser (GG) correction was used when the sphericity assumption was violated. Beyond hypothesis testing, we carried out and reported a series of exploratory analyses outlined in the preregistration. All analyses were performed using R version 3.5.1 with RStudio version 1.1.463. The raw data and code used for the analysis (R Markdown/HTML files) are publicly available at https://github.com/Pawel-Motyka/SMCVR.

## 3. Results

### 3.1. Performance in the sensorimotor mastery tasks

In line with the assumption of the study design, and constituting a useful manipulation check, the ability to control the objects differed significantly between the sensorimotor conditions. In the moving target task, the average distance between the pointer and the target (an inverse measure of performance) was lowest in the congruent condition, and successively greater in the opposite, novel, and random conditions, respectively, *F*(3, 141) = 343.28, *p* < 0.001, ε = 0.71, η^2^ _*G*_ = 0.81 (GG-corrected, all post hoc Bonferroni-corrected pairwise comparisons, *p* < 0.001; Fig. 4a). Correspondingly, in the static target task, the average number of hits on the target was the highest in the congruent condition and consecutively lower in the opposite, novel, and random conditions, *F*(3, 141) = 408.26, *p* < 0.001, ε = 0.77, η^2^ _*G*_ = 0.83 (GG-corrected, all comparisons *p* < 0.001, except for the comparison between the novel and random condition, *p* = 0.21; Fig. 4b). Thus, in both tasks, performance levels tended to reflect the expected difficulty of condition-specific sensorimotor dependencies. In contrast to the salience of sensorimotor conditions, the mere visual characteristics of the objects were not found to play a role in determining one’s performance (moving target task: *F*(3, 141) = 0.03, *p* = 0.992, η^2^_*G*_ < 0.001; static target task: *F*(3, 141) = 0.005, *p* = 0.999, η^2^ _*G*_ < 0.001; Fig. S1).

**Figure 4.**
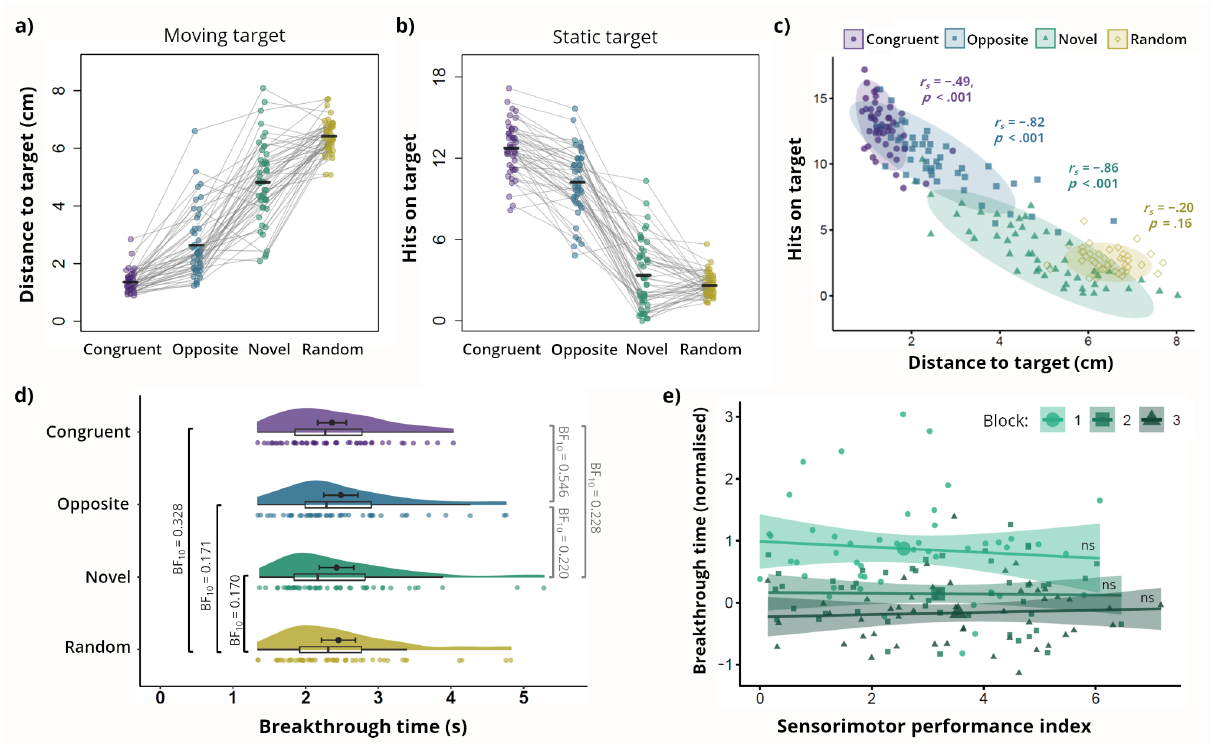
Summary of results. (**a, b**) In both sensorimotor mastery tasks, participants’ ability to control virtual objects decreased with the presumed difficulty of spatial coupling rules. **(c)** Performance measures from both tasks were moderately to strongly correlated (except for the random condition where no genuine learning was possible). **(d)** There were no significant differences in the average breakthrough times between objects manipulated under different sensorimotor conditions (BF_10_ < 0.333 indicate substantial evidence for the null hypothesis). Means are represented as thick points next to the boxplots (error bars show 95% confidence intervals). **(e)** No evidence for the acceleration of breakthrough times with an increasing ability to control virtual objects – neither in the most difficult but learnable novel condition (presented), nor in other conditions (see Figure S7).

To assess the learning of different sensorimotor dependencies over time, the changes in performance were analysed on a block-by-block basis. The analysis (ANOVA, within-subject factor: block) was run separately for different tasks and sensorimotor conditions. In the moving target task, a significant main effect of the block was observed only in the novel condition, *F*(2, 94) = 15.72, *p* < 0.001, ε = 0.91, η^2^_*G*_ = 0.06 (GG-corrected), where performance increased between the first two blocks (block 1-2, *t* = 3.90, *p* < 0.001; block 2-3, *t* = 1.53, *p* = 0.383; Fig. S2). In other conditions, there was no evidence for changes in performance over time (congruent: *F* = 0.45, *p* = 0.64, η^2^_*G*_ = 0.001; opposite: *F* = 1.08, *p* = 0.34, η^2^_*G*_ = 0.002; random: *F* = 1.27, *p* = 0.29, η^2^_*G*_ = 0.02). A similar pattern of results was observed in the static target task, in which the number of hits increased between all consecutive blocks only in the novel condition, *F*(2, 94) = 18.28, *p* < 0.001, ε = 0.93, η^2^_*G*_ = 0.05 (GG-corrected; block 1-2: *t* = −3.11, *p* = 0.007; block 2-3, *t* = −2.94, *p* = 0.013) and no significant effects were found under other conditions (congruent: *F* = 0.66, *p* = 0.52, η^2^ _*G*_ = 0.003; opposite: *F* = 1.92, *p* = 0.15, η^2^ _*G*_ = 0.004; random: *F* = 0.36, *p* = 0.70, η^2^ _*G*_ = 0.004; Fig. S3). To sum up, while it is unsurprising that there was no evidence of learning for the random and congruent conditions (in which respectively the floor- and ceiling-type effects could be expected), it is somewhat surprising that no improvement was found in the opposite condition. There, the performance level was relatively high and stable from the start – just as if one-to-one inversion was yet another example of a fairly familiar (or at least quickly learned) relationship. Evidence for ongoing learning was only found in the novel condition that relied on associations that are unlikely to occur in real-world situations (taking the form of a non-intuitive orthogonal mapping between the axes of action and those of visual rotation).

Next, we explored whether the velocity of performed manual movements varied across the sensorimotor conditions. In the moving target task, which involved tracking a shifting target, the average speed increased with the difficulty of the conditions, *F*(3, 141) = 102.72, *p* < 0.001, ε = 0.77, η^2^ _*G*_ = 0.36 (GG-corrected, congruent < opposite < novel < random; all comparisons *p* < 0.003). A slightly different pattern was observed in the static target task, *F*(3, 141) = 67.19, *p* < 0.001, ε = 0.59, η^2^ _*G*_ = 0.29 (GG-corrected), in which there was a nonsignificant trend for the lower average speed of movements at the opposite as compared to the congruent condition (*p* = 0.082; all other comparisons *p* < 0.002; opposite < congruent < novel < random). In general, the velocity was significantly higher in the task that required tracking of a continually shifting target (*M* = 1.23 degrees/frame) than in the task based on reaching a stationary target (*M* = 0.99 degrees/frame), *t*(47) = 10.94, *p* < 0.001. Additionally, we examined the associations between interindividual differences in movement speed and sensorimotor control performance across tasks and conditions. In the moving target task, increasing speed was associated negatively with performance (i.e., positively with distance to the target: congruent: *r*_*s*_ = 0.29, *p* = 0.044; opposite: *r*_*s*_ = 0.40, *p* = 0.005; novel: *r* = 0.31, *p* = 0.035; random: *r* = 0.39, *p* = 0.006). It is worth noting that in this task, the ideal speed would be that of the target itself, as in principle any excessive movement would result in moving away from the to-be-followed trajectory. In the static target task, however, action speed tended to be positively correlated with the number of hits in the relatively simpler conditions (congruent: *r* = 0.52, *p* < 0.001; opposite: *r*_*s*_ = 0.19, *p* = 0.186), while it was negatively correlated with performance in the novel condition which required adaptation to unfamiliar sensorimotor coupling rule (novel: *r*_*s*_ = −0.33, *p* = 0.023). This implies that once participants figured out how to rotate objects toward the target (particularly in simpler conditions), faster actions could only increase the total number of hits. As expected, movement speed was not significantly related to performance in the random condition (*r*_*s*_ = −0.01, *p* = 0.963), where target tracking seemed equally unfeasible regardless of action speed.

Finally, we examined the relationship between performance measures from both newly designed tasks. A moderate correlation was observed in the case of the congruent condition, *r*_*s*_ = −0.49, *p* < 0.001, and strong correlations were found for the opposite, *r*_*s*_ = −0.82, *p* < 0.001, and novel conditions, *r*_*s*_ = −0.86, *p* < 0.001 (Fig. 4c: negative correlations were expected due to opposite direction of the distance and hits measures). There was no significant correlation in the random condition, *r*_*s*_ = −0.20, *p* = 0.16, under which no genuine learning was possible. These results suggest that performance in both tasks was underpinned by the same ability to gain voluntary control over the direction of the object’s rotation, supporting the construct validity of these tasks and measures used. As planned in the preregistration, in the case of a correlation between the distance and hit measures, “a single index (e.g., the average of ranks or standardised values from both tasks)” was calculated and used in further analyses focused on whether the sensorimotor performance was associated with a decrease of breakthrough times in the CFS task. Such an index was derived by ranking the trial-level performances of all participants separately for each task and then averaging the task-specific ranks for each participant and condition. The ranking method was favoured as it seemed more insensitive to the impact of outlier observations. For convenience, the obtained sensorimotor performance index has been normalised to a value range of 0–10 (for more details, see the R Markdown file no. 4).

### 3.2. Breakthrough times of objects manipulated under different sensorimotor conditions

We hypothesised that objects manipulated under lawful sensorimotor dependencies (congruent, opposite, and novel conditions) would show faster access to visual awareness as compared to objects moving randomly in response to actions. However, we found no evidence for such an effect, *F*(3, 141) = 1.07, *p* = 0.37, η^2^ _*G*_ = 0.003, with the average breakthrough times being comparable between all sensorimotor conditions (Fig. 4d). To inform whether this nonsignificant result reflects genuine null effects rather than experimental insensitivity an additional Bayes factor analysis was performed (using the default Jeffreys–Zellner–Siow prior, as we lacked justification for informed prior specification). The results yielded substantial evidence for no differences (i.e., BF_10_ < 0.333) in breakthrough times across all three comparisons of interest (congruent vs random: BF_10_ = 0.328; opposite vs random BF_10_ = 0.171; novel vs random: BF_10_ = 0.170). Null-favouring evidence was found also in exploratory comparisons (congruent vs novel: BF_10_ = 0.228; opposite vs novel: BF_10_ = 0.220), except for the comparison between the congruent and opposite conditions where only a trend in support of the null hypothesis was observed (BF_10_ = 0.546). These results suggest that the employed manipulation of spatial sensorimotor dependencies was not a salient enough factor to affect detection times during the CFS.

In line with the preregistered plan, an analogous analysis was carried out using data from the last block only – where the most pronounced effects could potentially be observed. Again, there were no significant differences in breakthrough times between the sensorimotor conditions, *F*(3, 141) = 1.06, *p* = 0.378, η^2^_*G*_ = 0.005. Another analysis focusing on block-by-block changes in detection times (a two-by-two repeated measures ANOVA, factors: condition and block) yielded a significant main effect of the block, *F*(2, 94) = 40.07, *p* < 0.001, ε = 0.78, η^2^_*G*_ = 0.080 (GG-corrected), a nonsignificant effect of the condition, *F*(3, 141) = 0.71, *p* = 0.546, η^2^ _*G*_ = 0.002, and a nonsignificant interaction, *F*(6, 282) = 0.580, *p* = 0.746, η^2^_*G*_ < 0.001. For simplification, the effects of time passage were visualised and analysed separately for different conditions (Fig. S4). In all conditions, there was a significant main effect of the block (congruent: *F*(2, 94) = 20.78, *p* < 0.001, ε = 0.69, η^2^ _*G*_ = 0.110; opposite: *F*(2, 94) = 18.71, *p* < 0.001, ε = 0.98, η^2^_*G*_ = 0.070; novel: *F*(2, 94) = 33.98, *p* < 0.001, ε = 0.76, η^2^ _*G*_ = 0.090; random: *F*(2, 94) = 25.00, *p* < 0.001, ε = 0.79, η^2^_*G*_ = 0.060, all GG-corrected) with post hoc comparisons indicating a typically observed decrease of detection times with the duration of the experiment (Mastropasqua et al., 2015; Paffen et al., 2018).

Finally, we compared the average breakthrough times of different visual objects (regardless of their assignment to particular sensorimotor conditions). While an analogous analysis based on the pilot data did not yield significant effects (see section *Stimuli*), here we observed robust differences between almost all compared objects, *F*(3, 141) = 45.95, *p* < 0.001, ε = 0.86, η^2^ _*G*_ = 0.080 (GG-corrected; see Fig. S5: obj. 2 < obj. 1 < obj. 3 < obj. 4; with only one nonsignificant comparison between obj. 3 and obj. 4, *p =* 0.125; all other comparisons *p* < 0.001). These results are consistent with evidence that even stimuli similar in terms of low-level visual features (e.g., contrast and size) can exhibit differential access to visual awareness, likely due to inherent coarse-level differences in their visual characteristics (e.g., Gayet et al., 2019) that make stimuli distinguishable from one another in the first place. Despite standardisation efforts to equalise “baseline” detection times for different objects, it was presumed that with increasing statistical power these differences could become significant, while still allowing us to address the key question of this study with the use of a fully counterbalanced design.

In sum, in contrast to the evident effects of visual features on breakthrough times, the present findings did not yield evidence for the role of pre-exposure to different spatial sensorimotor dependencies in determining breakthrough times. Therefore, we found no evidence to support the hypothesis that exposure to different (but always contingent) sensorimotor spatial dependencies affects visual awareness.

### 3.3. Summary of detection and identification responses during the CFS

Conscious detection of an object was reported in 97.6% of all trials (7350/7488; the average percentage of stimuli detected per participant was 97.6%, *SD* = 4.66%, range: 80.8%–100%). Among trials with reported detection, 98.3% of the presented stimuli were correctly identified in the following 4-AFC task (2nd response). According to the preregistered plan, trials with reported detection but incorrect identification (126 trials, 1.70% of trials with detection) were excluded from all analyses except those reported in the current section (the average percentage of misidentifications per participant was 1.73%, *SD* = 1.86%, range: 0%–7.69%). Given the small number of such trials, the analysis exploring for which sensorimotor conditions and objects errors were more likely to occur was included only in the R Markdown file (no. 3). Next, we explored how likely participants were to correctly identify the object in trials where no detection was reported (2.4% of all trials). The results indicated that participants were able to do so significantly above the 25% chance level with an average accuracy of 49.7% (*SD* = 36.60%), *χ*^*2*^(1) = 59.7, *p* < 0.001. While this result could possibly be interpreted in terms of having a genuine ability to make correct guesses about sensory stimuli despite the lack of conscious perception (Mudrik et al., 2013; Vieira et al., 2017; for a broader discussion see: Kouider & Faivre, 2017; Stein, 2019; Stockart et al., 2024), it cannot be ruled that, at least in some cases, participants were able to notice the object (or its fragments) at the very end of the presentation. A closer exploration of how correct guesses might have depended on specific objects and conditions has also been included in the R Markdown file only (no. 3). Finally, we also analysed whether response times in the 4-AFC task differed between different conditions and objects. While there were no significant differences accounted for by the sensorimotor dependencies under which the objects were manipulated, *F*(3, 141) = 0.087, *p* = 0.967, η^2^_*G*_ < 0.001, identification times differed between the particular objects, *F*(3, 141) = 44.39, *p* < 0.001, ε = 0.85, η^2^ _*G*_ = 0.160 (GG-corrected, Fig. S6), with object 1 being indicated significantly faster than all others (potentially due to its use as an exemplary object in the instructions to the task).

### 3.4. Associations between performance in the sensorimotor mastery tasks and breakthrough times

Here we aimed to explore whether the acquisition of “sensorimotor mastery” was associated with the acceleration of perceptual access to visual objects during visual suppression. Our particular focus was on the novel condition in which actual block-by-block learning effects and large interindividual differences were observed. For the purposes of this analysis, breakthrough times were normalised for each object participant using the median absolute deviation (Leys et al., 2013). Condition-specific indexes of sensorimotor performance were then correlated with the corresponding breakthrough times. In none of the conditions was there a significant relationship between the measures from the sensorimotor mastery tasks and breakthrough times in the CFS task (congruent: *r*_*s*_ = −0.18, *p* = 0.226; opposite: *r* = 0.11, *p* = 0.476; novel: *r* = −0.03, *p* = 0.862; random: *r* = 0.20, *p* = 0.183). A complementary analysis, which relied on raw (i.e., not normalised) breakthrough times, also showed no significant correlations – neither when using the overall sensorimotor performance index nor distinct measures from static and moving target tasks (for more details, see the R Markdown file no. 4). Next, as the relationship between motor control and visual awareness could potentially only become apparent after prolonged exposure to different sensorimotor dependencies, further block-by-block analyses were performed. Again, the emphasis was placed on the novel condition, in which it would be particularly informative to observe that the relationship between sensorimotor performance and breakthrough times surfaces only in the later blocks or that it becomes stronger as the experiment continues. However, there was no significant relationship between these measures in any of the blocks of the novel condition (block 1: *r*_*s*_ = −0.08, *p* = 0.594; block 2: *r* = −0.02, *p* = 0.871; block 3: *r* = 0.06, *p* = 0.672; Fig. 4e). The same was observed for all other conditions analysed on a block-by-block basis (Fig. S7, all tests *p* > 0.112).

## 4. Discussion

Using the CFS paradigm embedded in immersive virtual reality, we investigated whether gaining practical knowledge of sensorimotor spatial dependencies governing interactions with virtual 3D objects enhances their access to visual awareness. In newly designed sensorimotor mastery tasks, we manipulated the complexity of the spatial mapping between the direction of manual movements and concomitant rotation of the objects. We hypothesised that stimuli whose motion was experienced as depending on the participant’s actions in a learnable manner (i.e., according to a congruent, opposite, or novel dependency) would break through suppression, in a continuous flash suppression paradigm, faster than stimuli moving randomly in response to actions (i.e., lacking world-related statistical structure to learn). While performance in the sensorimotor mastery tasks consistently decreased with the difficulty of the conditions (i.e., congruent > opposite > novel > random), the preregistered analysis yielded no significant differences in breakthrough times for objects using different coupling rules. An exploratory analysis which aimed to assess whether the acquisition of sensorimotor mastery was associated with accelerated conscious access to visual stimuli also showed no significant effects. Exploratory Bayesian analyses confirmed these conclusions. In summary, under the conditions used in this study, we found no evidence that knowledge of how an object responds to one’s actions plays a role in determining access to visual awareness.

The main novelty of the present work, compared to previous research using interocular suppression paradigms, lies in the use of goal-oriented tasks that allow the assessment of sensorimotor skills in a naturalistic 3D environment. We observed the expected condition-wise differences in sensorimotor performance as well as pronounced interindividual variation, including different learning rates for the most unfamiliar yet controllable ‘novel’ condition. In addition, the performance measures from two distinct tasks (based on reaching a stationary target or following a continuously changing rotation) were moderately to strongly correlated, supporting the overall validity of this approach. Moreover, in contrast to prior studies, we separated the part involving the active control of objects from the CFS part involving their perceptual suppression. This was done to rule out possible influences of the speed of visual movement, known to affect the detection of suppressed stimuli (Baker & Graf, 2008; Blake et al., 1998; Wade et al., 1984), and to more directly address the role of acquired knowledge about possible interactions with objects, thereby linking to the topic of affordances (Borghi & Riggio, 2015; Dalgarno & Lee, 2010; Gibson, 1979). Importantly, the role of ‘passive’ sensorimotor knowledge in shaping visual awareness remains debatable. For instance, Korisky & Mudrik (2021) reported enhanced perceptual access to familiar 3D objects compared to their 2D photographs and scrambled (i.e., ‘meaningless’) 3D-printed versions, interpreting these results in terms of preferential processing of objects that elicit plans for possible functional use. While there are multiple differences between their study and ours, the present results suggest that knowledge of specific *spatial* dependencies (i.e., being able to predict how objects will respond to actions) may have minimal to no effect on visual awareness. This implies that other explanations for the effects found by Korisky and Mudrik should also be considered, such as differences in visual familiarity between normal and scrambled 3D objects or more abstract (i.e., not necessarily action-related) knowledge of their meaning and functional use.

The present null findings could also be related to earlier research involving manual movements performed during and immediately before perceptual suppression, which showed no significant differences in perceptual access to stimuli rotating congruently or opposite to lifelong expectations (Suzuki et al., 2019; Dogge et al., 2018). This aligns with our theoretical assumption that these dependencies may be relatively easy to master, as evidenced by the high-performance levels in sensorimotor control tasks for both congruent and opposite conditions. Yet, a key extension compared to prior studies was the use of conditions where it was not possible (or much more difficult) to identify the spatial relationship between actions and their visual consequences. Even so, the expected increase in breakthrough times for randomly responding (as well as hard-to-control) objects was not observed, which adds to the growing evidence suggesting a general ineffectiveness of sensorimotor spatial manipulations in interocular suppression paradigms (Dogge et al., 2018; Motyka et al., 2021b; Paris et al., 2017; Suzuki et al., 2019).

Having said all this, it is noteworthy that – irrespective of the direction of action-induced changes – in all conditions, object movement remained *contingent* on the participant’s actions and such stimuli have already been shown to have prioritised access to visual awareness as compared to temporally uncoupled/pre-recorded inputs (Maruya et al., 2007; Suzuki et al., 2019; see also Motyka et al., 2021a). Therefore, a possible explanation of the present results is that mere sensorimotor contingency, by signalling the presence of some form of temporally close action–perception coupling, was salient enough to override any influences attributable to the identifiability of specific spatial dependencies. An alternative speculative interpretation is that the effects of sensorimotor mastery acquired with one group of objects might have transferred to other objects during the CFS task. Although we used distinct objects for each sensorimotor condition, their resemblance as “sphere-like objects” may have led to generalisation across conditions (e.g., that all objects might seem controllable when interacted with). This interpretation could be further assessed in a between-subjects study where only one type of sensorimotor dependency would be assigned to a given group. On top of this, since in our case sensorimotor coupling occurred prior to and not during perceptual suppression, comparisons with earlier reports should be treated with caution, as it is uncertain whether the outcomes would have differed had the procedure addressed ongoing visuomotor integration processes rather than knowledge of dependencies between actions and consciously observed visual changes.

Our hypothesis were in part motivated by the theory that varying levels of counterfactual richness of the models that encode the expected behaviour of particular objects should affect aspects of visual experience, such as ‘perceptual presence’ or ‘objecthood’ (Seth, 2014a). However, the present null findings do not carry clear implications for the theory itself and are in no position to empirically undermine it. First, it is important to note that a perceptual breakthrough during continuous flash suppression does not equate to perceptual presence, and operationalizing perceptual presence directly remains a challenge. Another reason is that it has not yet been determined how different aspects of bodily interactions with objects relate to feelings of perceptual presence (for a comprehensive discussion of this term and related concepts, see Seth, 2015; Wiese, 2015; Suzuki et al., 2023; Barkasi, 2021). Given our focus on manipulating sensorimotor spatial dependencies, other aspects of the sensorimotor environment were preserved in a manner typical of real-world conditions and virtual reality setups. In addition to the contingency between manual actions and object rotation, this included both temporally and spatially consistent changes in retinal stimulation linked to eye movements and small-scale head rotations. Thus, participants may still have retained the feeling that “more of the object could be uncovered through the right eye, head, or body movements” (Barkasi, 2021, p. 2545). In other words, it is possible that the sum of veridically preserved dependencies loaded the sense of presence to such an extent that the exact specificity of the spatial coupling had no further impact on its resultant level of presence (Suzuki et al., 2023). Thus, while we may have succeeded in creating highly realistic conditions of modulated affordances, the stimuli used did not fully mirror the features of the more fringe phenomena lacking objecthood or realness (e.g., synaesthetic concurrents, visual snow or afterimages), for which the underlying counterfactual richness is thought to be contrastingly poor due to a multifaceted decoupling from one’s movements (Seth, 2014a). While the development of theoretical models alone can guide the design of future manipulations and measures in this area, at least in the context of interocular suppression paradigms, we are gaining a clearer understanding of which aspects of bodily interaction with objects drive or fail to drive their prioritization in visual awareness.

In conclusion, this study used an immersive and interactive VR environment to examine the impact of sensorimotor spatial dependencies of different levels of complexity on conscious access to 3D objects. Despite clear differences in the ability to control objects governed by rules of increasing complexity, no corresponding effects on breakthrough times were found. This extends previous evidence suggesting the negligible role of spatial congruence (compared to contingency) between motor actions and their visual consequences in shaping perceptual experience under interocular suppression paradigms.

## Author contributions

PM: Conceptualization; Data curation; Formal analysis; Funding acquisition; Investigation; Methodology; Project administration; Resources; Software; Validation; Visualization; Writing - original draft; and Writing - review & editing.

DS: Conceptualization; Methodology; Resources; Supervision; Validation; Writing - review & editing.

AKS: Conceptualization; Methodology, Supervision; Validation; Writing - review & editing.

KS: Conceptualization; Methodology; Resources; Software; Supervision; Validation; Writing - review & editing.

## References

Alais, D., & Blake, R. (2005). Binocular Rivalry. MIT Press.

Auvray, M., Hanneton, S., & O’Regan, J. K. (2007). Learning to perceive with a visuo-auditory substitution system: Localisation and object recognition with “the vOICe.” Perception, 36(3), 416–430. 10.1068/p5631

Baker, D. H., & Graf, E. W. (2008). Equivalence of physical and perceived speed in binocular rivalry. Journal of Vision, 8, 26–26. 10.1167/8.4.26

Barkasi, M. (2021). Does what we dream feel present? Two varieties of presence and implications for measuring presence in VR. Synthese, 199(1), 2525–2551. 10.1007/s11229-020-02898-4

Blake, R., Yu, K., Lokey, M., & Norman, H. (1998). Binocular rivalry and motion perception. Journal of Cognitive Neuroscience, 10, 46–60. 10.1162/089892998563770

Bermejo, F., Di Paolo, E. A., Hüg, M. X., & Arias, C. (2015). Sensorimotor strategies for recognizing geometrical shapes: A comparative study with different sensory substitution devices. Frontiers in Psychology, 6. https://www.frontiersin.org/article/10.3389/fpsyg.2015.00679

Borghi, A. M., & Riggio, L. (2015). Stable and variable affordances are both automatic and flexible. Frontiers in Human Neuroscience, 9, 351. 10.3389/fnhum.2015.00351

Clark, A. (2013). Whatever next? Predictive brains, situated agents, and the future of cognitive science. The Behavioral and Brain Sciences, 36(3), 181–204. 10.1017/S0140525X12000477

Dalgarno, B., & Lee, M. J. (2010). What are the learning affordances of 3-D virtual environments? British Journal of Educational Technology, 41(1), 10–32.

Degenaar, J. (2014). Through the inverting glass: First-person observations on spatial vision and imagery. Phenomenology and the Cognitive Sciences, 13(2), 373–393. 10.1007/s11097-013-9305-3

Dogge, M., Gayet, S., Custers, R., & Aarts, H. (2018). The influence of action-effect anticipation on bistable perception: Differences between onset rivalry and ambiguous motion. Neuroscience of Consciousness, 2018(1). 10.1093/nc/niy004

Gayet, S., & Stein, T. (2017). Between-Subject Variability in the Breaking Continuous Flash Suppression Paradigm: Potential Causes, Consequences, and Solutions. Frontiers in Psychology, 8. https://www.frontiersin.org/article/10.3389/fpsyg.2017.00437

Gayet, S., Paffen, C. L. E., Belopolsky, A. V., Theeuwes, J., & Van der Stigchel, S. (2016). Visual input signaling threat gains preferential access to awareness in a breaking continuous flash suppression paradigm. Cognition, 149, 77–83. 10.1016/j.cognition.2016.01.009

Gayet, S., Stein, T., & Peelen, M. V. (2019). The danger of interpreting detection differences between image categories: A brief comment on “Mind the snake: Fear detection relies on low spatial frequencies” (Gomes, Soares, Silva, & Silva, 2018). Emotion, 19, 928–932. 10.1037/emo0000550

Gibson, J. J. (1979). The theory of affordances. The ecological approach to visual perception. Boston: Houghton Mifflin.

Held, R. (1965). Plasticity in sensory-motor systems. Scientific American, 213(5), 84–94. 10.1038/scientificamerican1165-84

Hense, M., Badde, S., & Röder, B. (2019). Tactile motion biases visual motion perception in binocular rivalry. Attention, Perception & Psychophysics, 81(5), 1715–1724. 10.3758/s13414-019-01692-w

Hohwy, J. (2013). The predictive mind. Oxford University Press.

Hohwy, J., Roepstorff, A., & Friston, K. (2008). Predictive coding explains binocular rivalry: An epistemological review. Cognition, 108(3), 687–701. 10.1016/j.cognition.2008.05.010

Kaspar, K., König, S., Schwandt, J., & König, P. (2014). The experience of new sensorimotor contingencies by sensory augmentation. Consciousness and Cognition, 28(100), 47–63. 10.1016/j.concog.2014.06.006

Kohler, I. (1951 [1964]). The formation and transformation of the perceptual world (trans. H. Fiss). Psychol. Issues 3(4), Monograph 12.

Korisky, U., & Mudrik, L. (2021). Dimensions of Perception: 3D Real-Life Objects Are More Readily Detected Than Their 2D Images. Psychological Science, 32(10), 1636–1648. 10.1177/09567976211010718

Kouider, S., & Faivre, N. (2017). Conscious and Unconscious Perception. In The Blackwell Companion to Consciousness (pp. 551–561). John Wiley & Sons, Ltd. 10.1002/9781119132363.ch39

Leys, C., Ley, C., Klein, O., Bernard, P., & Licata, L. (2013). Detecting outliers: Do not use standard deviation around the mean, use absolute deviation around the median. Journal of Experimental Social Psychology, 49(4), 764–766. 10.1016/j.jesp.2013.03.013

Lunghi, C., Morrone, M. C., & Alais, D. (2014). Auditory and Tactile Signals Combine to Influence Vision during Binocular Rivalry. Journal of Neuroscience, 34(3), 784–792. 10.1523/JNEUROSCI.2732-13.2014

Maruya, K., Yang, E., & Blake, R. (2007). Voluntary Action Influences Visual Competition. Psychological Science, 18(12), 1090–1098. 10.1111/j.1467-9280.2007.02030.x

Mastropasqua, T., Tse, P. U., & Turatto, M. (2015). Learning of monocular information facilitates breakthrough to awareness during interocular suppression. Attention, Perception & Psychophysics, 77(3), 790–803. 10.3758/s13414-015-0839-z

Miles, W. R. (1930). Ocular Dominance in Human Adults. The Journal of General Psychology, 3(3), 412–430.

Motyka, P., Kozłowska, Z., & Litwin, P. (2021a). Perceptual Awareness of Optic Flows Paced Optimally and Non-optimally to Walking Speed. Perception, 50(9), 797–818. 10.1177/03010066211034368

Motyka, P., Akbal, M., & Litwin, P. (2021b). Forward optic flow is prioritised in visual awareness independently of walking direction. PLOS ONE, 16(5), e0250905. 10.1371/journal.pone.0250905

Mudrik, L., Gelbard-Sagiv, H., Faivre, N., & Koch, C. (2013). Knowing where without knowing what: Partial awareness and high-level processing in continuous flash suppression. Journal of Vision, 13(9), 1103. 10.1167/13.9.1103

Nöe, A. (2004). Action in Perception. A Bradford Book.

O’Regan, J. K., & Noë, A. (2001). A sensorimotor account of vision and visual consciousness. The Behavioral and Brain Sciences, 24(5), 939–973; discussion 973-1031. 10.1017/s0140525x01000115

Paffen, C. L. E., Gayet, S., Heilbron, M., & Van der Stigchel, S. (2018). Attention-based perceptual learning does not affect access to awareness. Journal of Vision, 18(3), 7. 10.1167/18.3.7

Parker, A. L., & Alais, D. M. (2006). Auditory modulation of binocular rivalry. Journal of Vision, 6(6), 855–855. 10.1167/6.6.855

Paris, R., Bodenheimer, B., & Blake, R. (2017). Does direction of walking impact binocular rivalry between competing patterns of optic flow? Attention, Perception & Psychophysics, 79(4), 1182–1194. 10.3758/s13414-017-1299-4

Parr, T., Corcoran, A. W., Friston, K. J., & Hohwy, J. (2019). Perceptual awareness and active inference. Neuroscience of Consciousness, 2019(1), iz012. 10.1093/nc/niz012

Sachse, P., Beermann, U., Martini, M., Maran, T., Domeier, M., & Furtner, M. R. (2017). “The world is upside down” – The Innsbruck Goggle Experiments of Theodor Erismann (1883–1961) and Ivo Kohler (1915–1985). Cortex, 92, 222–232. 10.1016/j.cortex.2017.04.014

Schumann, F., & O’Regan, J. K. (2017). Sensory augmentation: Integration of an auditory compass signal into human perception of space. Scientific Reports, 7(1), 42197. 10.1038/srep42197

Seth, A. K. (2014a). A predictive processing theory of sensorimotor contingencies: Explaining the puzzle of perceptual presence and its absence in synesthesia. Cognitive Neuroscience, 5(2), 97–118. 10.1080/17588928.2013.877880

Seth, A. K. (2014b). The Cybernetic Bayesian Brain. In T. Metzinger & J. M. Windt (Eds.), Open MIND. Open MIND. Frankfurt am Main: MIND Group. 10.15502/9783958570108

Seth, A. K. (2015). Presence, objecthood, and the phenomenology of predictive perception. Cognitive Neuroscience, 6(2–3), 111–117. 10.1080/17588928.2015.1026888

Stein, T. (2019). The breaking continuous flash suppression paradigm: Review, evaluation, and outlook. In Transitions between consciousness and unconsciousness, 1st ed (pp. 1–38). Routledge/Taylor & Francis Group. 10.4324/9780429469688-1

Stockart, F., Schreiber, M., Amerio, P., Carmel, D., Cleeremans, A., Deouell, L., Dienes, Z., Elosegi, P., Gayet, S., Goldstein, A., Halchin, A.-M., Hesselmann, G., Kimchi, R., Lamy, D., Loued-Khenissi, L., Meyen, S., Micher, N., Pitts, M., Salomon, R., … Mudrik, L. (2024). Studying unconscious processing: Contention and consensu. 10.31234/osf.io/bkxzh

Suzuki, K., Schwartzman, D. J., Augusto, R., & Seth, A. K. (2019). Sensorimotor contingency modulates breakthrough of virtual 3D objects during a breaking continuous flash suppression paradigm. Cognition, 187, 95–107. 10.1016/j.cognition.2019.03.003

Suzuki, K., Mariola, A., Schwartzman, D. J., & Seth, A. K. (2023). Using Extended Reality to Study the Experience of Presence. Current Topics in Behavioral Neurosciences. 10.1007/7854_2022_401

Tsuchiya, N., & Koch, C. (2004). Continuous flash suppression. Journal of Vision, 4(8), 61. 10.1167/4.8.61

Wade, N. J., De Weert, C. M. M., & Swanston, M. T. (1984). Binocular rivalry with moving patterns. Perception & Psychophysics, 35, 111–122. 10.3758/BF03203891

Wähnert, S., & Gerhards, A. (2022). Sensorimotor adaptation in VR: Magnitude and persistence of the aftereffect increase with the number of interactions. Virtual Reality, 26(3), 1217–1225. 10.1007/s10055-022-00628-4

Wiese, W. (2014). Perceptual Presence in the Kuhnian-Popperian Bayesian Brain. In T. Metzinger & J. M. Windt (Eds.), Open MIND. Open MIND. Frankfurt am Main: MIND Group. 10.15502/9783958570207

Vieira, J. B., Wen, S., Oliver, L. D., & Mitchell, D. G. V. (2017). Enhanced conscious processing and blindsight-like detection of fear-conditioned stimuli under continuous flash suppression. Experimental Brain Research, 235(11), 3333–3344. 10.1007/s00221-017-5064-7

Veto, P., Schütz, I., & Einhäuser, W. (2018). Continuous flash suppression: Manual action affects eye movements but not the reported percept. Journal of Vision, 18(3), 8. 10.1167/18.3.8

Veto, P., Uhlig, M., Troje, N. F., & Einhäuser, W. (2018). Cognition modulates action-to-perception transfer in ambiguous perception. Journal of Vision, 18(8), 5. 10.1167/18.8.5

Yamamoto, K., Hoffman, D. S., & Strick, P. L. (2006). Rapid and long-lasting plasticity of input-output mapping. Journal of Neurophysiology, 96(5), 2797–2801. 10.1152/jn.00209.2006

Zopf, R., Schweinberger, S. R., & Rich, A. N. (2019). Limits on visual awareness of object targets in the context of other object category masks: Investigating bottlenecks in the continuous flash suppression paradigm with hand and tool stimuli. Journal of Vision, 19(5), 17. 10.1167/19.5.17

