## Supplemental material for "Investigating the role of sensorimotor spatial dependencies in shaping conscious access to virtual 3D objects"


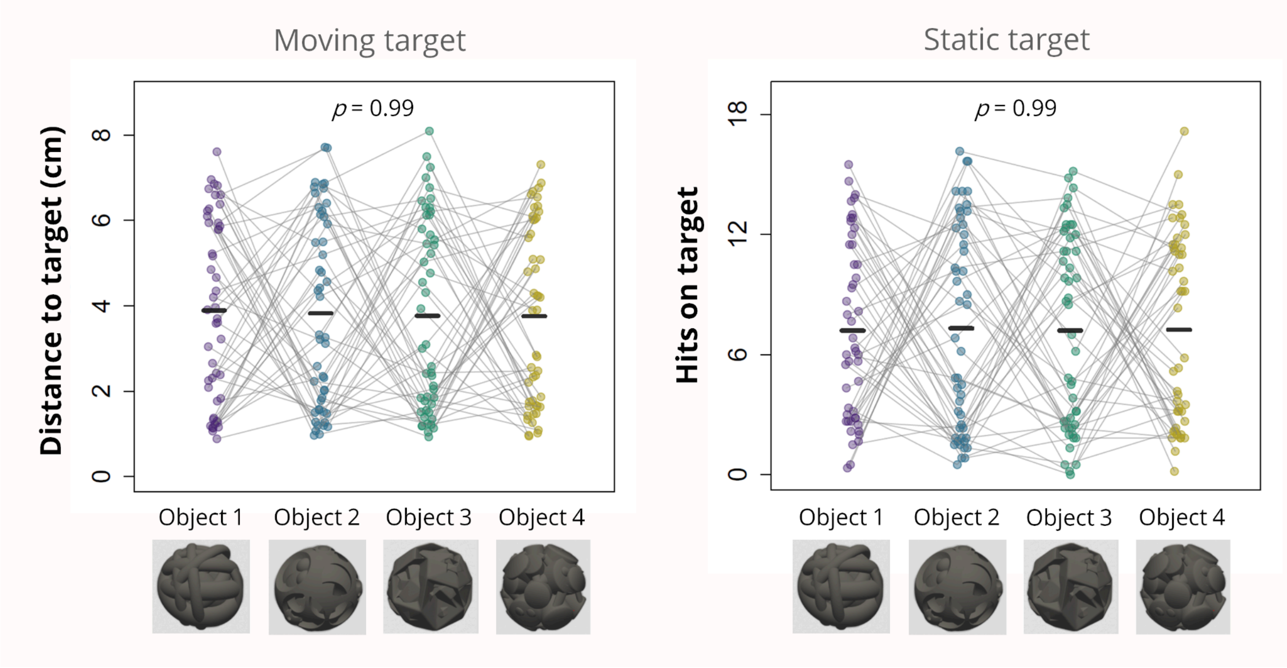


**Figure S1.** No significant differences in sensorimotor control over different visual objects.

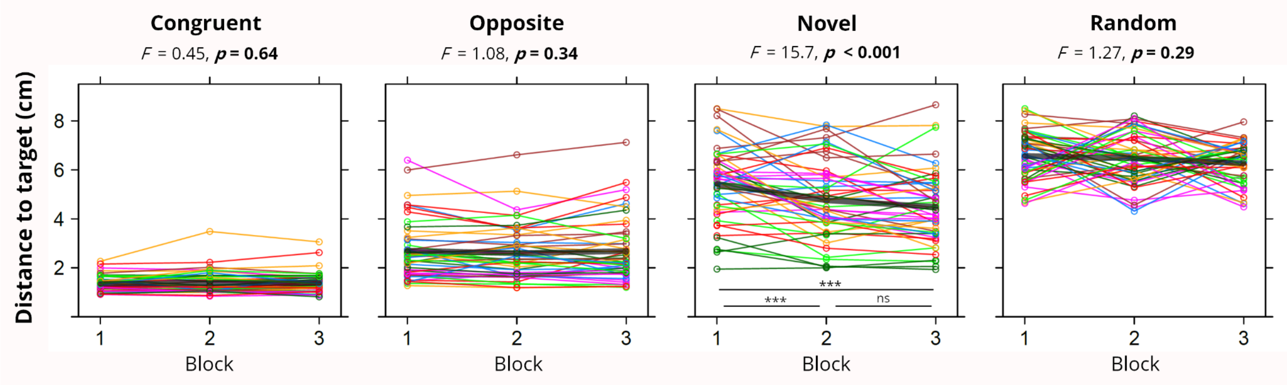


**Figure S2.** Block-by-block changes in sensorimotor performance under different conditions – moving target task.

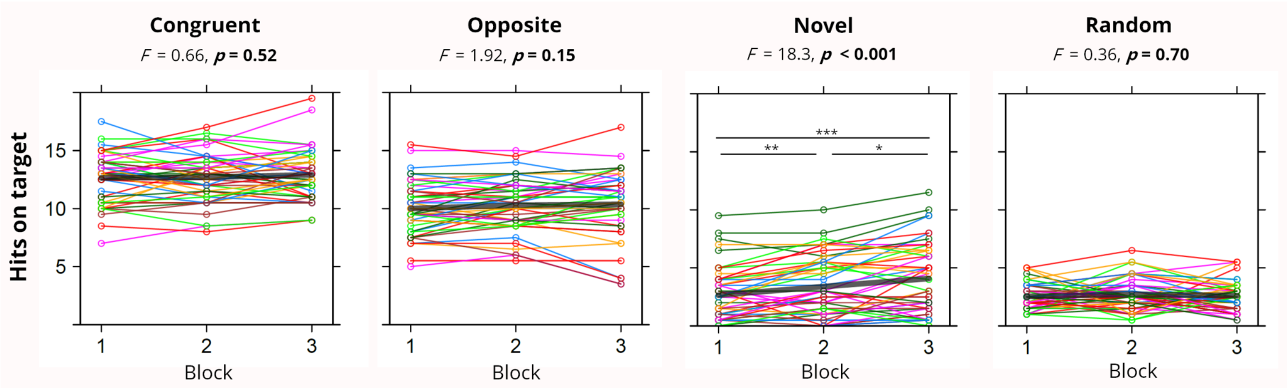


**Figure S3.** Block-by-block changes in sensorimotor performance under different conditions – moving target task.

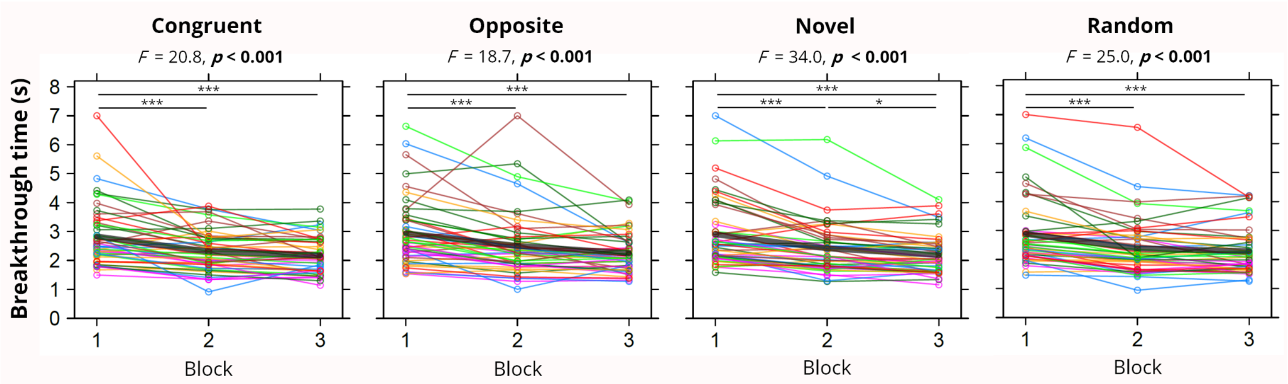


**Figure S4.** Block-by-block changes in breakthrough times under different conditions (CFS task).

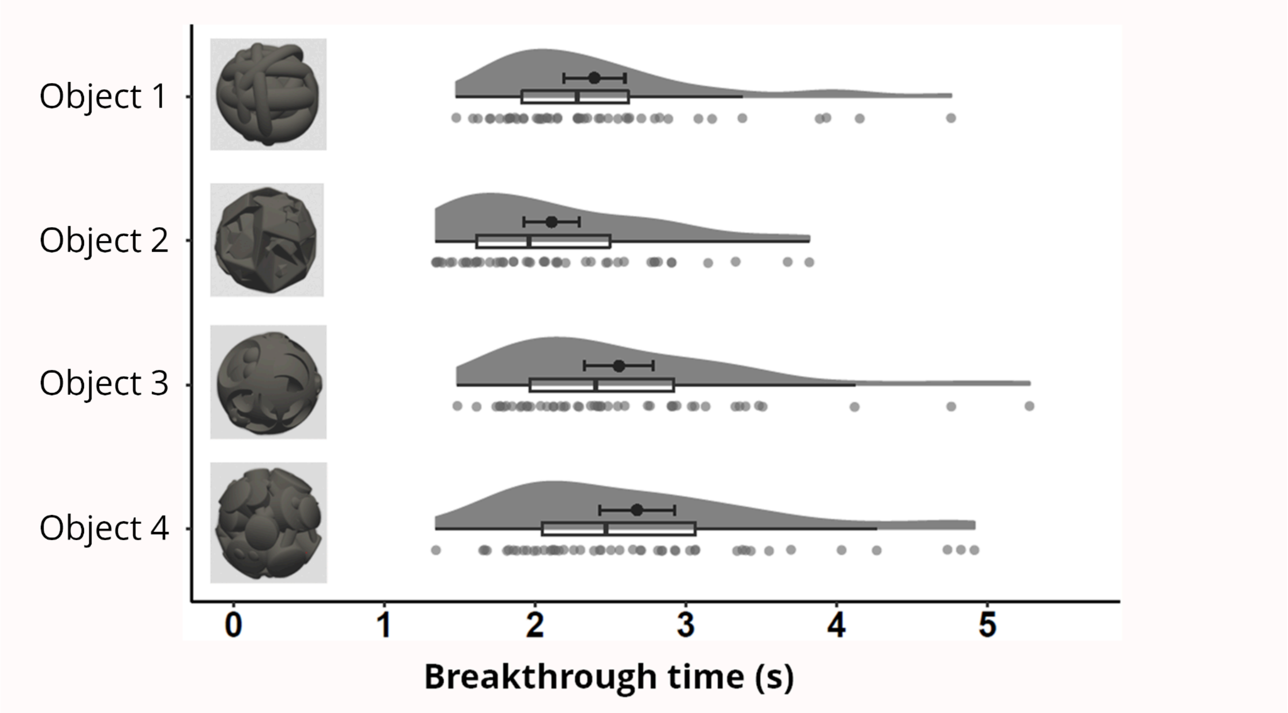


**Figure S5.** Average breakthrough times for different visual objects (CFS task). Means are represented as thick points next to the boxplots (error bars show 95% confidence intervals).

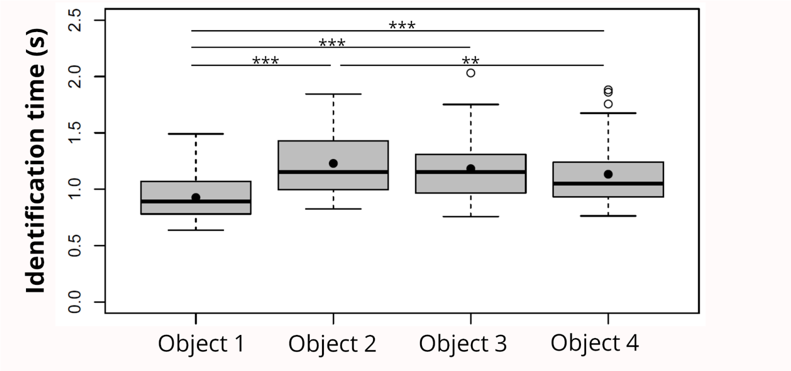


**Figure S6.** Average identification times for different objects (CFS: 4-AFC task).

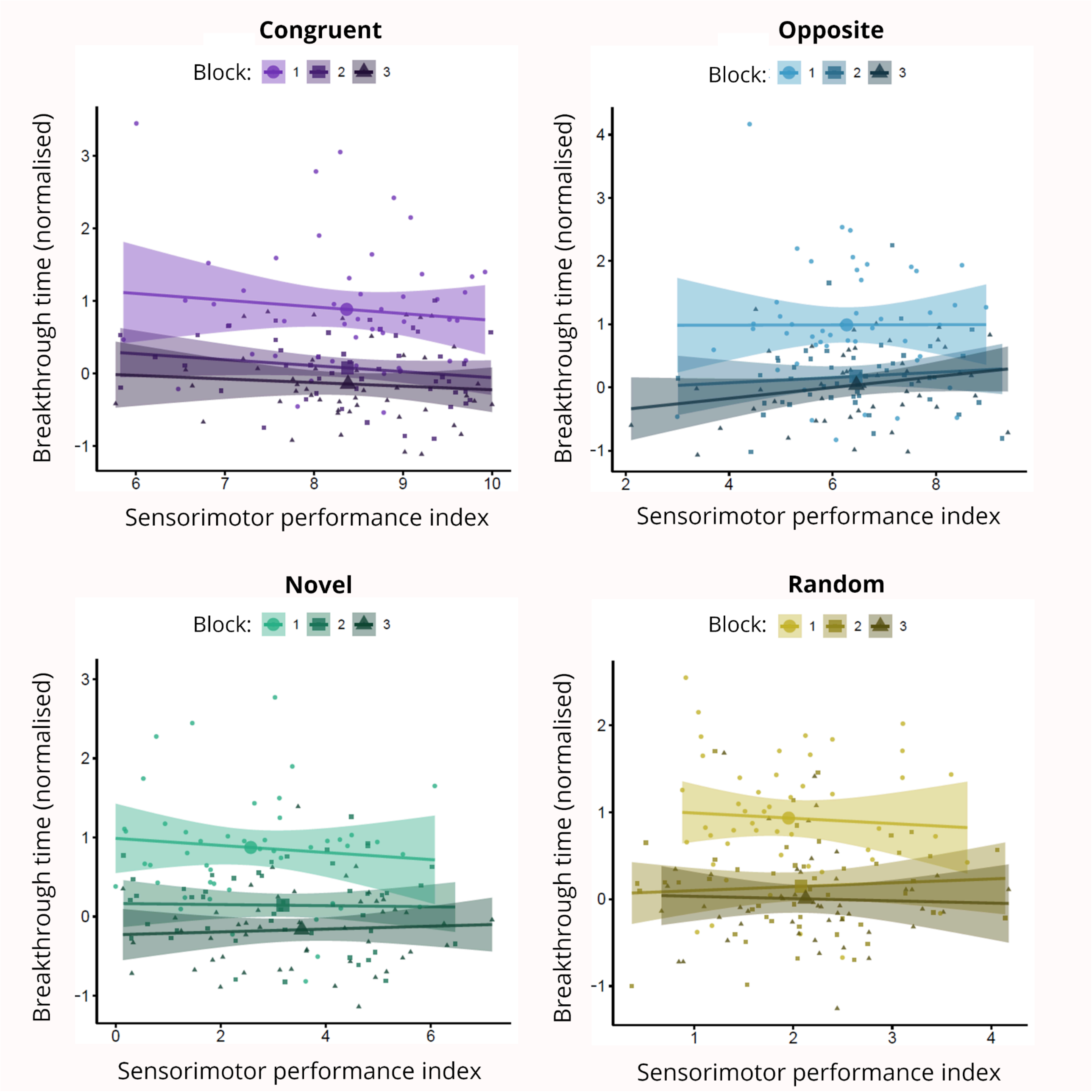


**Figure S7.** Relationships between sensorimotor performance index (x-axis) and the average normalised breakthrough time (y-axis) across different conditions.

Supplementary video 1: Static target task; https://www.youtube.com/watch?v=uZ0kcg1nnXM
Supplementary video 2: Moving target task; https://www.youtube.com/watch?v=3J_F7_Cw4jA
Supplementary video 3: CFS task; https://www.youtube.com/watch?v=6P2OM2tQvFw
